# AI discovery of sequence rules for RNA polymerase II pausing revises the pause-release model of gene activation

**DOI:** 10.64898/2026.09.18.752771

**Authors:** Nova Fong, Nicolas Huynh, Benjamin Erickson, Ryan M. Sheridan, Srinivas Ramachandran, Mihaela van der Schaar, David L. Bentley

## Abstract

The promoter-proximal pause (PPP), at most metazoan genes, is believed to prime transcriptional activation by release of paused RNAPII into the gene. We tested the pause-release model by abolishing the PPP and monitoring gene activation. An LLM-guided AI discovered that upstream G’s and a G T/C dinucleotide robustly discriminates pauses from non-pauses. Insertion of G-less sequences into NDRG1 and HSP90AA1 abolished the PPP in an orientation-dependent way, but did not impair activation by hypoxia or heat-shock. Hence pause-release is dispensable for these responses. The 5′ peak of RNAPII detected by ChIP persisted even when pausing, detected by NET-seq was absent. Instead of pausing, these promoter-proximal polymerases may undergo abortive initiation. RNAPII C-terminal domain (CTD) deletion suppresses the PPP and shifts it upstream without affecting sequence-dependent pausing, consistent with promoter-tethering by this IDR. We propose that PPP formation is initiated by G-dependent arrest, and stabilized by multiple factors including CTD-dependent tethering.

## Introduction

A signature feature of transcription in most metazoans is that RNA polymerase II pauses about 40-60 bases downstream of the start site. It was proposed over 30 years ago that pre-loading and promoter-proximal pausing of RNAPII ^1-7^ enables gene activation by release into the gene body in response to numerous signals including heat shock and hypoxia ^8-13^. Positive transcription elongation factor b, PTEFb comprising CDK9/CyclinT is required for RNAPII release from the PPP, and inhibition of this kinase by dichlororibofuranosylbenzimidazole (DRB) causes RNAPII arrest at the PPP ^14^ and inhibits elongation. Transcription elongation complexes are stabilized at the PPP by Negative Elongation Factor (NELF), and DRB Sensitivity Inducing Factor (DSIF, Spt4/5), that contact the polymerase and the RNA transcript ^15-19^. However depletion of neither NELF nor DSIF in vivo prevents PPP formation ^20,21^. DNA sequence motifs that bind TFIID and GAGA factor ^22,23-26^ correlate with the presence of a PPP but no sequence essential for its establishment has been identified. As a result it has never been possible to eliminate the PPP either generally or gene-specifically in order to test its function directly. High levels of turnover of RNAPII at the PPP ^27-29^ are mediated by premature termination factors including integrator and CRL3^ARMC5^ ^3,30-33^. In vitro, abortive initiation is another active mechanism for turnover of earlier transcription complexes that have not escaped the promoter ^34,35^ ^36^ but its importance in vivo is not known.

Mapping of pause sites in human cells by nascent elongating RNA sequencing (NET-seq) ^37,38^ showed that they are enriched at G,T/C dinucleotides with a minor preference for G’s within the transcription bubble ^21^, but other than this modest amount of sequence information, little is known about why pausing occurs at some positions and not others. The same consensus sequence applies to sites of pausing within the PPP where RNAPII piles up to high levels and within the gene body ^21^ where average RNAPII occupancy remains low. Why high peaks of RNAPII density only occur close to transcriptional start sites (TSS) is incompletely understood. One idea is that the RNAPII peak is positioned by the +1 nucleosome ^39^ but conversely paused RNAPII may help position the +1 nucleosome ^40,41^. Another possible explanation is that tethering of polymerase to the promoter helps to establish the PPP. The RNAPII C-terminal domain (CTD), which is a highly extended intrinsically disordered region (IDR) can make dynamic multivalent contacts with the IDR’s of transcription factors ^42,43^. It has been suggested that formation of bridging contacts between the CTD and the IDRs of promoter-bound transcription factors may help establish the PPP ^44^. The CTD comprises conserved heptad repeats that are reversibly phosphorylated at different positions in a manner that is synchronized with the transcription cycle ^45,46^. CTD Ser5 phosphorylation by TFIIH-associated CDK7/CyclinH occurs at initiation of transcription ^47-49^ and subsequent phosphorylation by PTEFb is associated with release from the PPP ^50^. CTD phosphorylation can control the dissolution of phase-separated condensates with other IDRs ^43,51^. Promoter-tethering by the CTD has yet to be experimentally tested, but a notable precedent occurs in *E. coli* where it is the basis of pausing that depends on promoter-bound σ70 ^52^.

## Results

### AI Discovery of DNA features that discriminate pause from non-pause sites

To identify sequence features that distinguish pause from non-pause sites, we applied Dynamic Engineering of Features in Trees (DEFT), a Large Language Model (LLM)-guided framework for growing interpretable decision trees ^53^. The tree was constructed based on 101-bp sequence windows centered on the candidate pause position, defined here as position +50, from approximately 95,000 pause sites and a matched set of non-pause sites (Fig. 1). Pause sites were selected at random from throughout the length of genes in HCT116 cells ^21^. DEFT does not operate as a black-box predictor. It embeds an LLM inside the learning algorithm and proposes at each tree node, human-readable sequence features, converts them into executable rules, and retains only those that improve discrimination between pause and non-pause sites. Each discovered feature is represented semantically, enabling prediction and interpretation. This approach differs fundamentally from other AI DNA analysis tools ^54-56^ that extract regulatory signals by interpreting predictors trained for predefined tasks. DEFT identified a composite sequence feature—G at the pause position, C or T at the +1 position, together with a minimum G content in the 49 bases upstream—as the primary discriminator between pause and non-pause sites. The high-G branch of this first split was strongly enriched for pausing, containing 66,897 pause sites but only 6,680 non-pause sites, corresponding to ∼69% of pause sites and ∼7% of non-pause sites in the dataset (Fig. 1). Conversely, the low-G branch was strongly enriched for non-pause sites. In particular, in this branch, sites with lower upstream G content contained 75,560 of the non-pause sites (Fig. 1). To assess whether the features identified by DEFT generalize beyond the training data, we evaluated the decision tree learned by DEFT on a held-out test set comprising 48,357 samples: 24,178 pauses and 24,179 non-pauses. On this held-out set, the model discriminated between pause and non-pause sites with an AUROC (area under the receiver operating characteristic curve) of 0.92 and an accuracy of 0.87 (Table S1).

**Figure 1.**
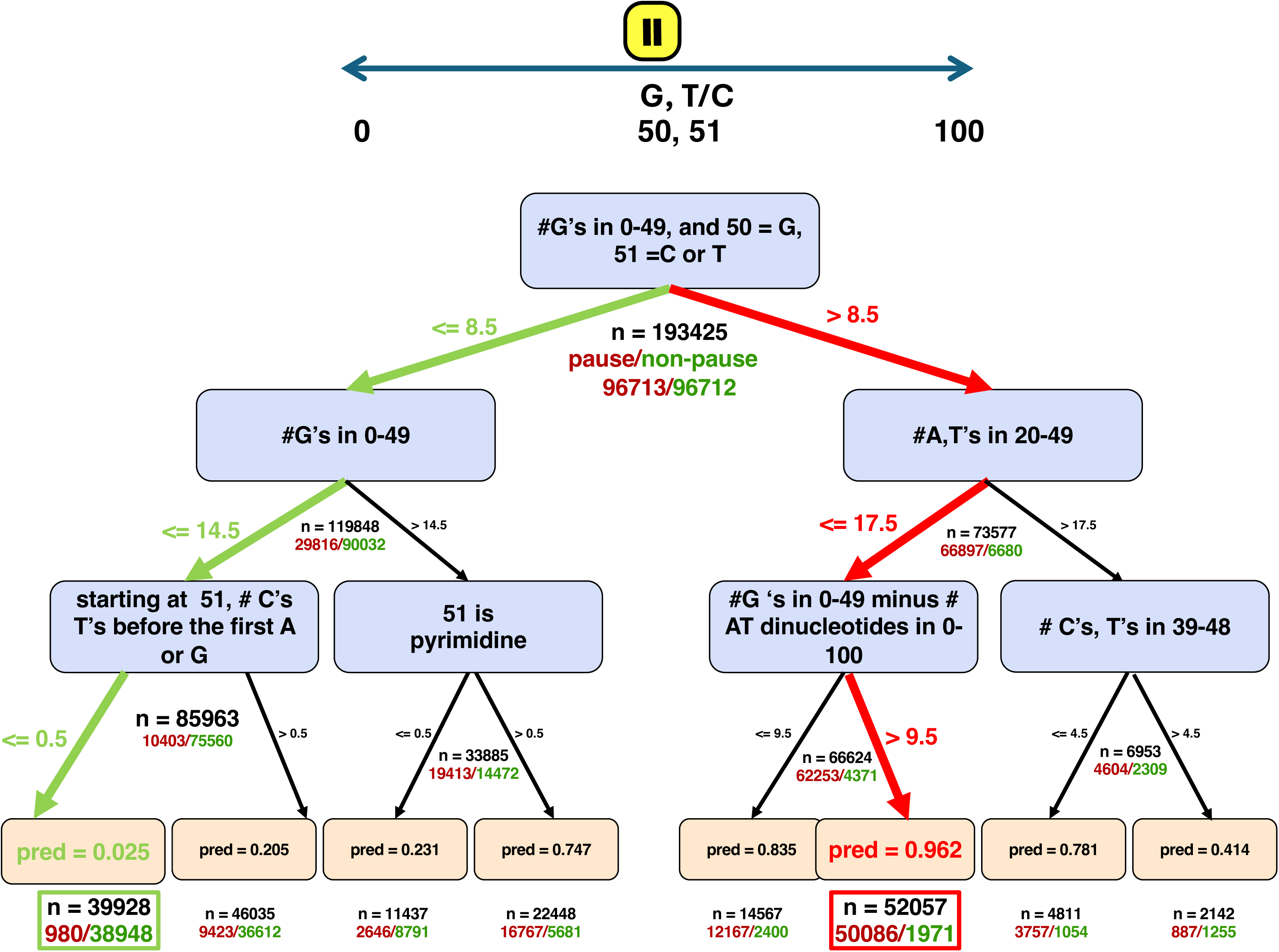
Decision tree analysis shows upstream G content predicts RNAPII pause sites. Pauses and non-pauses from eNETseq analysis of HCT116 cells ^21^(GEO: GSE202749 <u>GSM6132198</u>, <u>GSM6132200</u>). Sequence features identified by DEFT ^53^ within a 101 base region that are highly predictive of pausing (red) and non-pausing (green). The pause site at position 50 is enriched for G preceding a pyrimidine. Pauses are positions where eNETseq signal is 3 standard deviations above the mean for a window of 200 bases (100 bases upstream and downstream of the site) ^21,37^. Red and green arrows define paths that identify sequences highly enriched for pauses and non-pauses respectively.

### Abolition of the PPP by insertion of a G-less sequence

It is not known if pause sites within the PPP differ from those within the gene body but their consensus sequences are almost indistinguishable ^21^. We therefore presumed that features detected by DEFT might apply both in the PPP and the gene body. Based on the prediction that pausing requires a minimum upstream G content, we inserted a synthetic sequence lacking G’s at the 5’ end of the hypoxia inducible NDRG1 gene. A 377 bp G-less cassette was inserted by CRISPR Cas9 mediated homology directed repair into HAP1 cells at +8 relative to the transcription start site (Fig. 2A). This G-less cassette was developed to facilitate in vitro transcription assays ^57^ and is long enough to detect effects on pausing within the PPP and further downstream. The NDRG1 promoter has a TATA box and C_-1_A_+1_ initiation site but not a canonical initiator element ^58^. The insertion upstream of the first G of the transcript is not expected to disrupt promoter function since TATA containing promoters generally lack a downstream core promoter region ^59^. RNA-seq confirmed that the transcript starts at the beginning of the G-less sequence (Fig. 2B). Using eNET-seq ^21^, we mapped the 3’ ends of nascent transcripts immunoprecipitated with RNAPII +/-dimethyloxalylglycine (DMOG) that mimics hypoxia by stabilizing HIF1α ^60^. Reads were aligned to a custom genome that included the G-less insertion. Remarkably, the G-less sequence abolished the PPP in control and DMOG activated cells (Fig. 2C red arrow). As a control, the cassette was inserted at the same position in the inverted orientation i.e a C-less cassette (Fig. 2A). Inversion of the G-less cassette restored a prominent PPP 30 bases downstream of the TSS in two independent cell lines (Fig. 2D, blue arrow). These effects of the G-less and C-less cassette insertions were specific to sense transcription while divergent antisense transcription was unaffected (Fig. 2E, antisense strand). To control for the possibility of poor mapping to the G-less sequence, we mapped synthetic NETseq data matched for read-length distribution to the custom human genome and found no evidence of reduced reads at the G-less region (Fig. S1A). To check whether pause suppression was specific to HAP1 cells, we inserted the G-less cassette into both copies of NDRG1 in HCT116 cells and observed a similar abolition of the PPP (Fig. 2F, red arrow). The G-less insertion effectively shifts the initially transcribed sequence (ITS) where the PPP normally occurs, 380 bases downstream from the TSS. At this downstream position the ITS no longer supported a RNAPII peak - or + DMOG activation (Fig. 2C, F open blue arrows).

**Figure 2.**
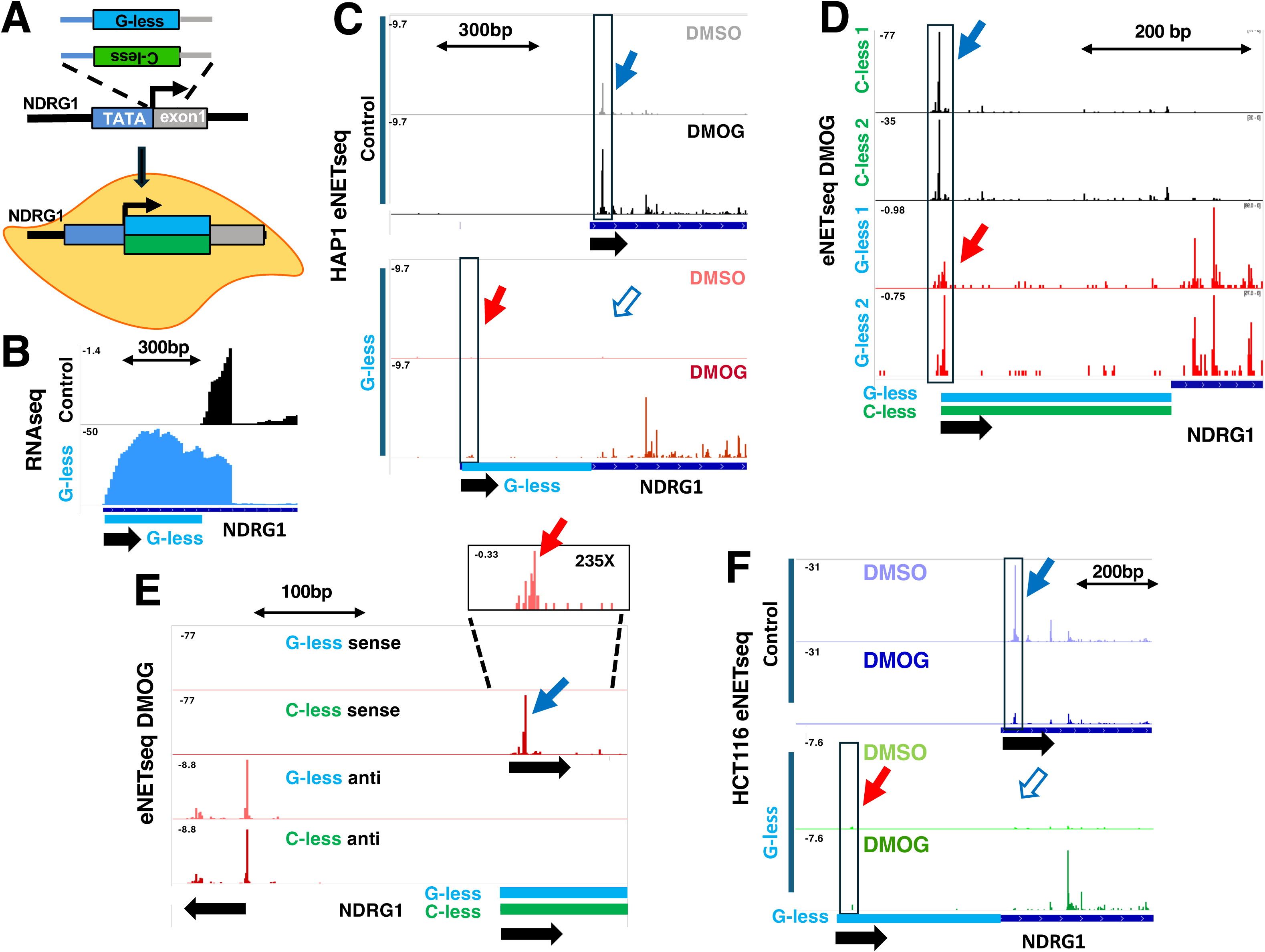
PPP suppression by G-less insertion shows that pause release is dispensable for hypoxia responsive gene activation. **A.** CRISPR insertion of G-less and inverted C-less cassettes (377bp) at the 5’ end of NDRG1. **B.** RNAseq of the 5’ end of DMOG activated NDRG1 with and without the G-less cassette insertion. **C.** eNETseq reads (3’ ends of nascent transcripts) at 5’ end of NDRG1 gene -/+ DMOG in control HAP1 cells (top panel) and cells with the G-less insertion (bottom panel). Note the PPP (solid blue arrow) is abolished by the G-less insertion (red, open blue arrow). **D.** eNETseq for two independent isolates of the G-less and inverted C-less cassettes at NDRG1 in HAP1 cells +DMOG. The PPP is lost with the G-less cassette (red arrow) and restored when it is inverted (C-less. blue arrow). Note the differences in Y axis scale. **E.** eNETseq reads as in B at 5’ end of NDRG1 with G-less or inverted C-less cassettes + DMOG. Sense (top two tracks) and divergent antisense (bottom two tracks) transcription from the NDRG1 promoter are shown. The inverted C-less insertion restores the PPP (blue arrow) by >100X relative to the G-less sequence (red arrow) whereas divergent antisense transcription is unaffected. **F.** eNETseq -/+DMOG in HCT116 cells with and without the G-less cassette at NDRG1. The PPP is suppressed by the G-less cassette (red arrow). Note differences in Y axis scale.

A small peak of NETseq signal was detected within the G-less cassette at the position of the PPP (∼ +40) in DMOG induced cells (Fig. 2E inset, red arrow), but it was >100 fold reduced relative to the inverted C-less control and far smaller than pauses within the gene body. The C-less insertion also demonstrates that a synthetic sequence can support a strong PPP in vivo. Pausing was also suppressed within the ∼300 bases of G-less sequence downstream of the PPP and was restored within the C-less cassette (Fig. 3A, blue arrows). Thus PPP suppression by the G-less region is orientation-dependent, consistent with an effect of sequence, rather than a biophysical property like its ability to localize nucleosomes. The G-less cassette is predicted to accommodate two nucleosomes with a small offset between the two orientations and similar localization scores as determined by NuPoP V2.5 ^61^(Fig. S1B). The strong orientation-dependent suppression of the PPP is therefore unlikely to be caused by a difference in nucleosome occupancy.

**Figure 3.**
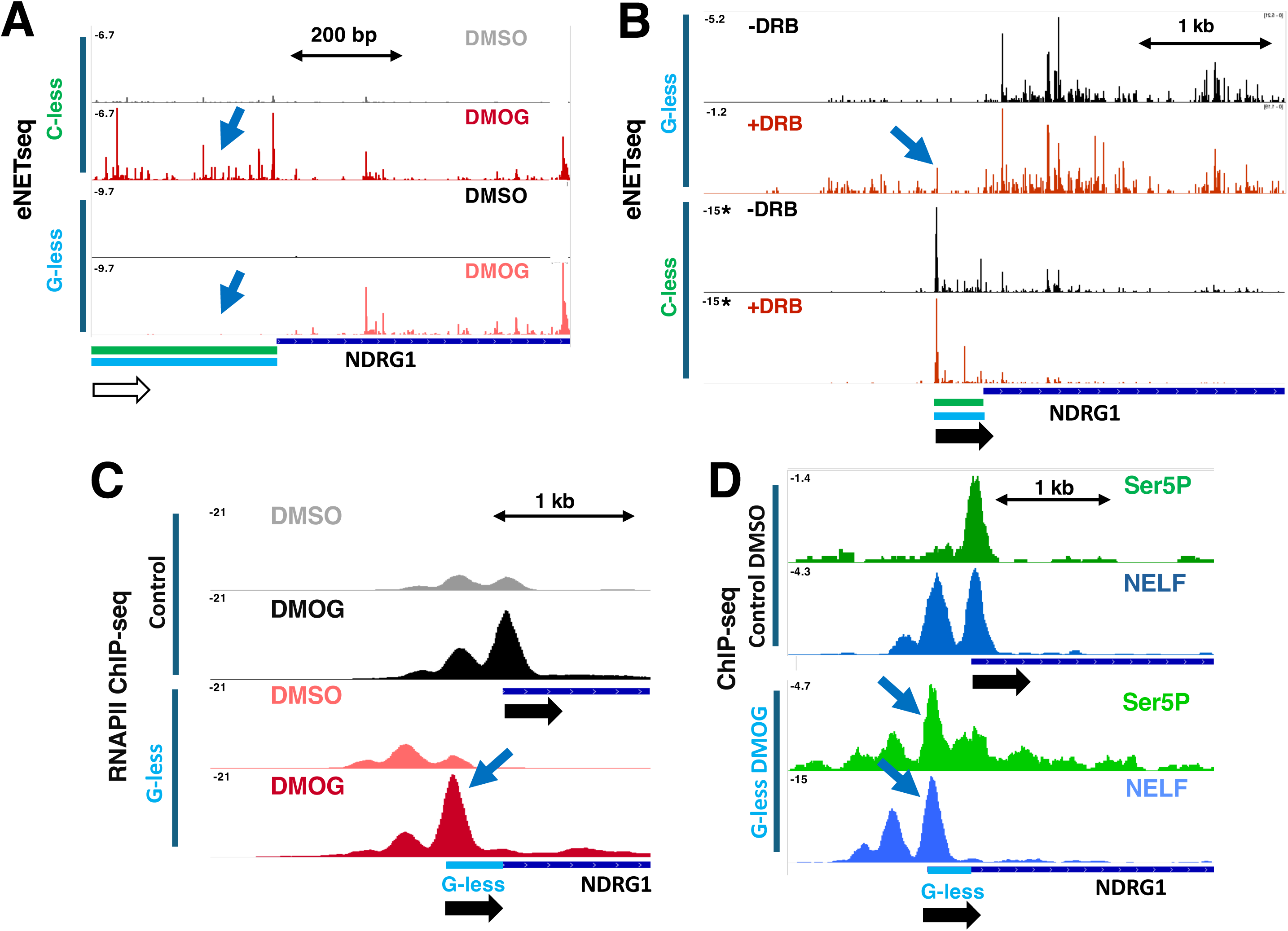
A 5’ peak of RNAPII and NELF in the absence of pausing. **A.** eNETseq showing suppression of pausing within the G-less region relative to the C-less region + DMOG (blue arrows). Note the region shown is downstream of the TSS and PPP. **B.** eNETseq of NRDRG1 with G-less and inverted C-less insertions + DMOG +/-DRB (100 μM, 2 hr). Note DRB does not restore a PPP to the gene with the G-less cassette. * full range of the Y axis not shown. **C.** ChIP-seq shows promoter-proximal peak of RNAPII (blue arrow) is unaffected by the G-less cassette which suppresses pausing as detected by NET-seq (Fig.2). **D.** Promoter proximal peaks of Ser5-P RNAPII and NELF (blue arrows) occur at the G-less gene in the absence of a promoter-proximal pausing.

We asked whether inhibition of PTEFb with DRB could rescue the PPP at a G-less sequence. As expected DRB limited elongation on control genes (Fig, S1C). Although DRB limited elongation within the DMOG-activated NDRG1 gene downstream of a G-less sequence, it did not restore a PPP at the start site (Fig. 3B, S1D, blue arrows). Thus G-dependent pausing is necessary for PPP establishment even when elongation is impaired by inhibiting PTEFb. We conclude that, as predicted by DEFT, upstream G residues on the non-template strand are essential for formation of the PPP and for pausing downstream of the PPP in the gene body.

### The 5’ peak of RNAPII is not necessarily paused

We compared the 5’ RNAPII peaks detected by eNETseq versus anti-RNAPII ChIP-seq. eNET-seq maps the 3’ ends of nascent transcripts at least 22-25 bases long on fully engaged elongation complexes^37^ whereas ChIP-seq detects any RNAPII close enough to the template to be X-linked by formaldehyde. Surprisingly, while the G-less sequence almost totally suppresses pausing detected by eNETseq, it has little effect on total RNAPII occupancy at the 5’ end of the gene as detected by ChIP (Fig. 3C). As expected CTD Ser5 phosphorylation that accompanies transcription initiation was also detected by ChIP-seq at the 5’ end of the inserted G-less cassette (Fig. 3D, blue arrow). The pausing factor NELF co-localized with the RNAPII ChIP peak at the 5’ end of the G-less sequence although there is no detectable pausing (Fig. 3D). The disparity between pausing detected by NET-seq and RNAPII occupancy detected by ChIP indicates that while the G-less sequence prevents pausing of elongation complexes, it does not reduce RNAPII recruitment or CTD Ser5 phosphorylation associated with initiation. We speculate that RNAPII that piles up in the absence of pausing is associated with RNA transcripts too short to be captured by NETseq. Such complexes may undergo abortive initiation which is very frequent in vitro ^34-36^. These results therefore show that, contrary to what is commonly thought, the 5’ peak of RNAPII ChIP signal does not necessarily equate with pausing and some conclusions based on this assumption may need to be revised.

### Pause-release is dispensable for gene activation at NDRG1 and HSP90AA1

The pause-release model, predicts that preventing PPP establishment would impair gene activation. We asked how PPP suppression by a G-less sequence affects transcription, measured as RNAPII density within the gene body, before and after activation of NDRG1. The G-less insertion did not detectably de-repress basal transcription under control (DMSO) conditions however it substantially increased the fold activation by DMOG as measured by eNETseq or RNAPII ChIPseq (Fig. 4A, B, S1E). As expected, activation of other hypoxia inducible genes was unaffected (Fig. S1F). These observations suggest that suppression of the PPP with a G-less ITS not only fails to inhibit transcriptional activation of a hypoxia inducible gene, but can actually enhance it. Higher RNAPII occupancy in the gene body downstream of the G-less cassette following DMOG stimulation was not associated with increased strength or frequency of pauses (Fig. 4C) suggesting that it is not the result of slower transcription, nor was there an obvious loss of processivity across the 65kb long gene (Fig. 4A, B, S1E). Together these findings suggest that PPP suppression by the G-less sequence elevates transcriptional flux through the gene body. This conclusion is consistent with RNAseq of NDRG1 mRNA which revealed a super-activation in response to DMOG because of the G-less cassette insertion (Fig. 4D, S1G). Similarly RT-PCR analysis showed that NDRG1 mRNA activation by DMOG or CoCl_2_ was enhanced by the G-less cassette insertion but not by the C-less insertion (Figs. 4E, S2A). Notably the RNAseq profile for NDRG1 does not reveal any evidence of exon skipping, intron retention or defective 3’ end formation associated with the G-less cassette insertion (Fig. 4D).

**Figure 4.**
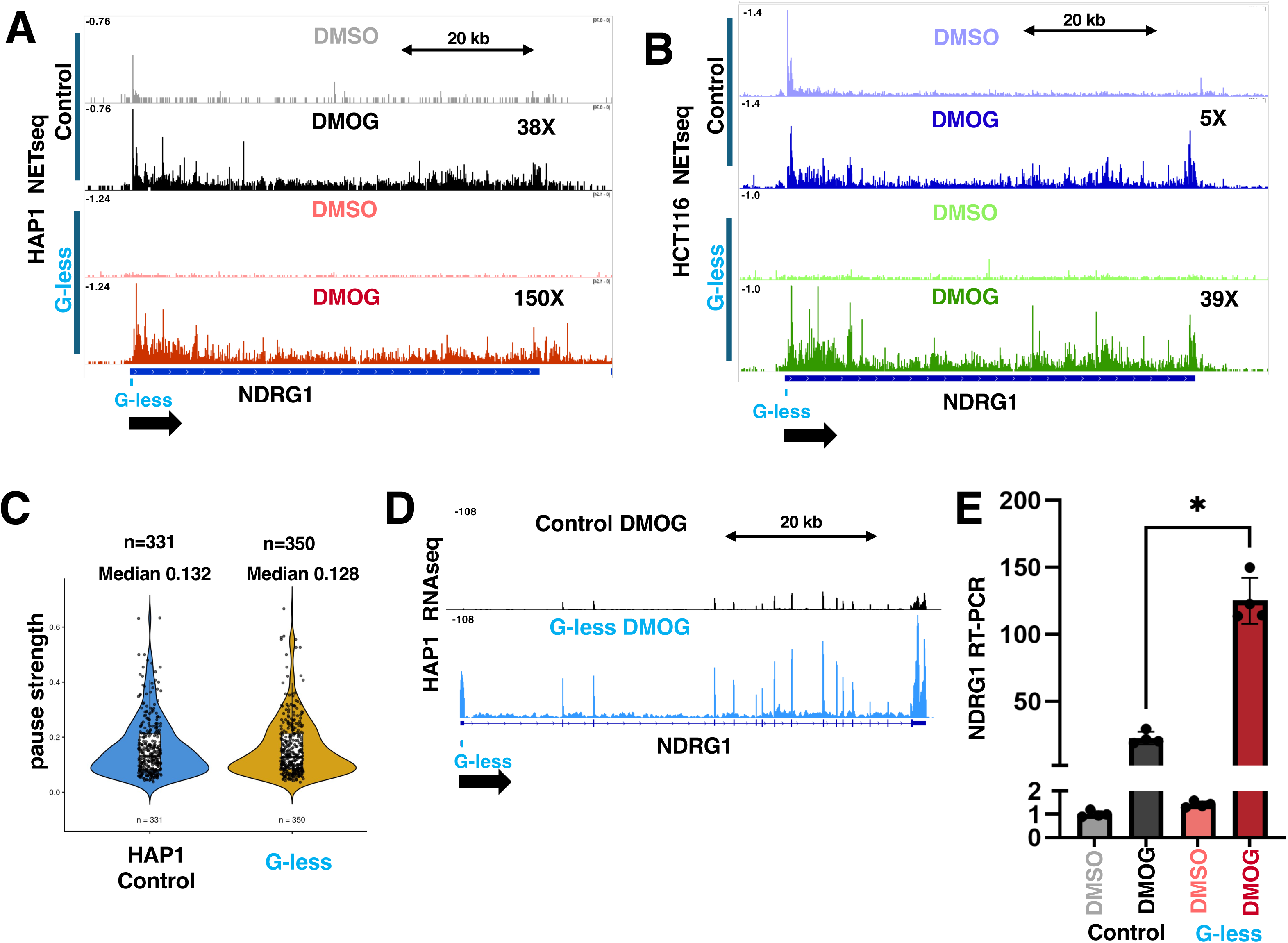
The PPP is dispensable for transcriptional activation of NDRG1. **A, B.** eNETseq +/-DMOG in HAP1 and HCT116 with and without G-less cassette insertion. Fold activation determined at the ratio of gene body signal +/-DMOG. Note super-activation when pausing is suppressed by the G-less insertion, **C.** Pause strength in the NDRG1 gene body in HAP1 defined as (# reads at pause) / (# reads in 200bp window) after subsampling to equalize gene body read numbers. G-less insertion does not significantly increase pause number or strength**. D.** RNAseq of DMOG activated NDRG1 with and without the G-less cassette insertion. Note super-activation by the G-less insertion (see Fig. S1G for ACTB control). **E.** RT-PCR of NDRG1 mRNA induction by DMOG shows super-activation in HAP1 cells with the G-less cassette insertion. We did not observe super-activation of NDRG1 mRNA by the G-less insertion in HCT116, possibly due to post-transcriptional regulation.

To investigate whether G’s in the ITS are required for PPP formation at other genes, we inserted the G-less cassette at the TSS of HSP90AA1 which has a TATA box promoter and a C_-1_A_+1_ initiator. The insertion was made at +9 and G’s at +2 and +5 were changed to A which is not expected to disrupt to the promoter significantly^58^. eNETseq showed that, the G-less ITS at HSP90AA1 abolished the PPP at 37° without affecting divergent antisense transcription (Fig. 5A, S2B, red arrows) as we observed at NDRG1. Also consistent with the NDRG1 G-less gene, Ser5P RNAPII and NELF were recruited to the 5’ end of the G-less HSP90AA1 gene (Fig. S2C, blue arrows). Despite abolition of the PPP at HSP90AA1 (but not other heat shock genes, Fig. S2D), rapid transcriptional activation in response to heat shock (30 min. 42°) was relatively unaffected by the G-less insertion (Fig. 5B, S2E). We did not observe a consistent super-activation of HSP90AA1 with the G-less cassette insertion as we did for NDRG1. We conclude that at both hypoxia and heat shock inducible genes, a G-less ITS effectively abolishes the PPP without impairing gene activation.

**Figure 5.**
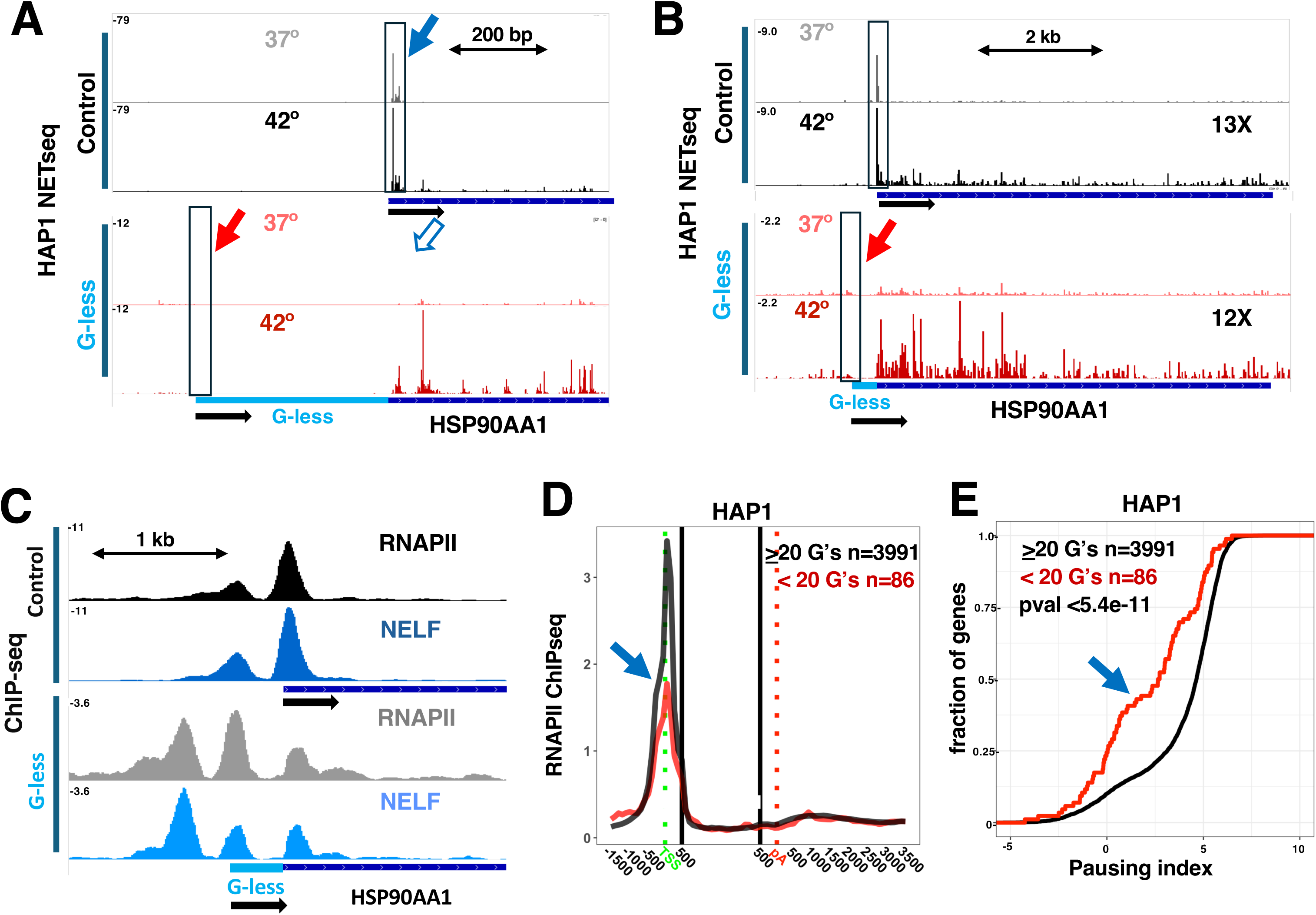
Rapid heat shock activation in the absence of a PPP. **A.** eNETseq at HSP90AA1 at 37° and 42° (30 min) in HAP1 -/+ G-less cassette insertion at 37° and 42°. Note the PPP present in control cells (blue arrow) is abolished by the G-less insertion (open blue and red arrows) **B.** eNETseq as in A showing fold increase of eNETseq reads in the gene body 42°/37°. **C.** Promoter proximal peaks of RNAPII and NELF HSP90AA1 (37°) occur at the G-less insertion gene (bottom two tracks) in the absence of pausing. **D.** RNAPII ChIPseq in HAP1 for genes in the top 50% for expression with <20 or >20 G’s in the first 100 bases of the transcript. Note reduced PPP (blue arrow) for low G content genes. **E.** Cumulative frequency plots of pausing index (PI=Log2 (RNAPII density -30 - +300/+301 - polyA) for genes with low and high G content as in D. Note that low G content is associated with significantly reduced PPP (blue arrow).

### Reduced PPP at genes with low 5’ G content

We investigated the PPP at human genes with low G content at their 5’ ends. G is the most abundant base at the 5’ ends of human genes with a median count of 36 in the first 100 bases (Fig. S2F). Genes with fewer than 20 G’s in the first 100 bases are rare among the top 50% of expressed human genes (Fig. S2F), and they have a significantly reduced PPP and lower pausing index than other genes, consistent with the importance of G’s for PPP establishment (Fig. 5D, E, Fig. S3A, B). HoxB2 is one such gene that has only 14G’s and 60 A’s+T’s in the first 100 bases of the transcript and lacks a detectable PPP (Fig. S3C). In future it will be of interest to determine the minimum number of G’s and their distribution within the ITS that is necessary for PPP formation.

### The RNAPII CTD affects the strength and position of the PPP

While G-dependent pausing is necessary for PPP establishment, it is clearly not sufficient since pausing occurs at the same G, T/C motifs within the gene body ^21^ (Fig. 6A, S3G, left panels) where large peaks of RNAPII density do not build up. In vitro, NELF and DSIF stabilize the PPP ^16-18^ ^15^ but in vivo depletion of these factors has little effect on the PPP ^20,21^ suggesting that additional factors contribute. We asked whether the CTD might contribute to promoter tethering of paused polymerases, as recently proposed ^44^, by performing eNETseq on RNAPII with epitope tagged full-length large subunit, Rpb1, or Rpb1ΔCTD (Figure 6B). This experiment showed that the principal effect of CTD deletion is to suppress the PPP and reduce the pausing index at 1000’s of genes (Fig. 6C, S3E) including immediate early (e.g. FOS, ARC), heat shock (DNAJA1, HSPA1B) and hypoxia (LDHA) responsive genes (Fig. 6D, S4). This is unlikely to be an indirect effect of altered NELF or DSIF association since these factors do not bind the CTD^62^. Notably, CTD deletion appears to mimic CTD phosphorylation that also promotes early elongation ^50,63,64^ and is predicted to disrupt IDR-IDR interactions ^43,44,51^. We enquired whether reduced promoter proximal pausing by RNAPII ΔCTD resulted from poor recognition of G, T/C pauses, but found no evidence for less pausing at these sites at either promoter proximal or distal positions (Fig. 6A, Fig. S3G right panels). The CTD can adopt conformations up to ∼50 nm long ^65^, that could span over 100 bases of extended DNA. A prediction of the tethering model is that CTD deletion would prevent pausing at positions more distal from the TSS. Remarkably this prediction was confirmed by eNETseq which showed that the peak of paused RNAPII shifts from ∼ +80 bases relative to the TSS for WT to ∼ +40 for ΔCTD RNAPII (Fig. 6E, S3F). In summary the CTD stimulates PPP establishment or maintenance independent of G-specified pausing. In addition the effect of the CTD on the position of the PPP is consistent with the idea that it functions as a tethering arm that retains RNAPII at a distance from contact points in the promoter as previously suggested ^44^.

**Figure 6.**
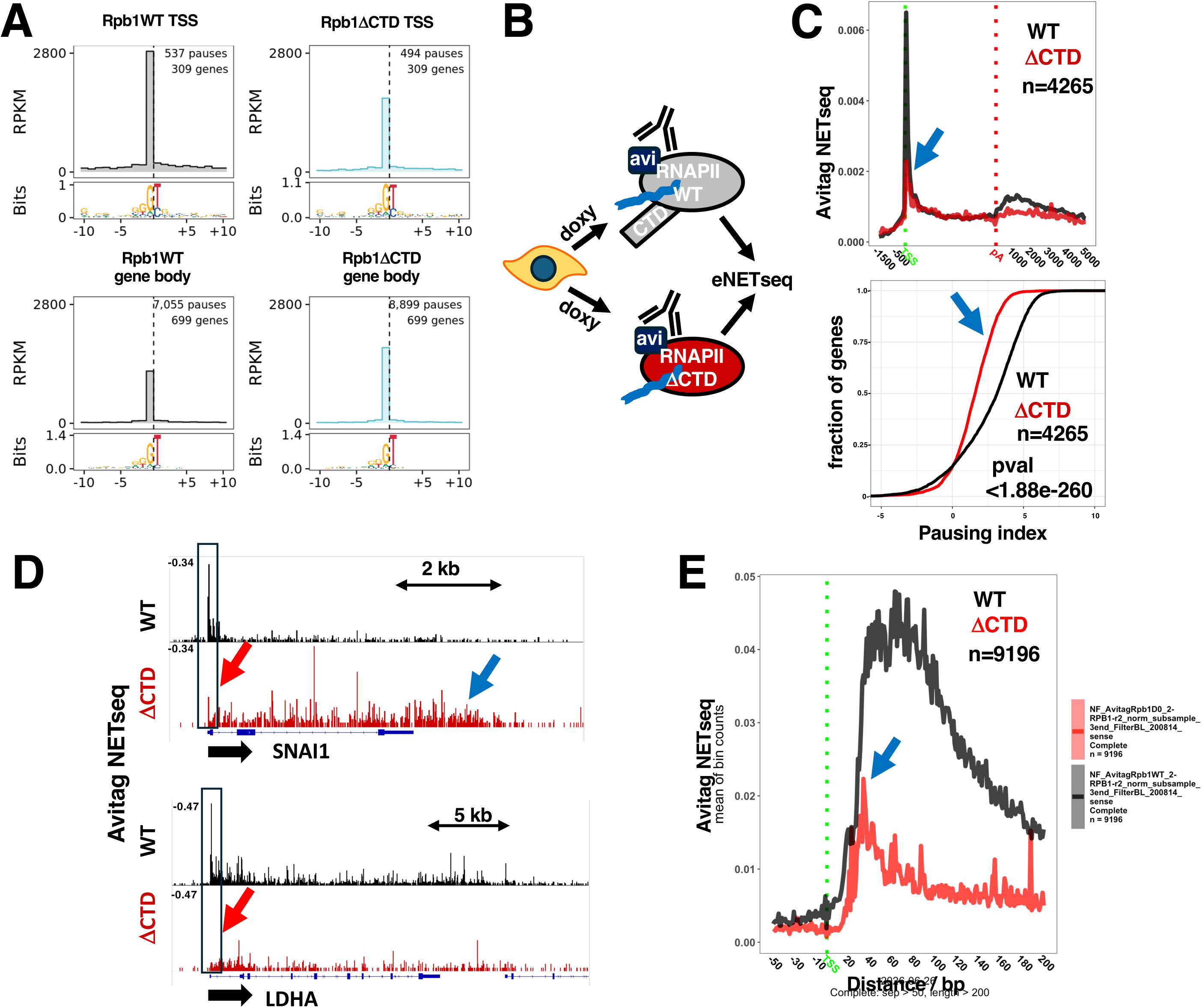
The RNAPII CTD enhances promoter-proximal pausing. **A.** Consensus sequences around pause sites (dotted line) for WT and ΔCTD RNAPII. Pauses were identified after down-sampling to equalize reads in the two samples within each region: TSS(+1 - +500 bp) and gene body (+501-polyA). ΔCTD does not reduce pausing at G, T/C consensus sites. **B.** Strategy for eNETseq of RNAPII WT and ΔCTD using doxycycline inducible mutants of Rpb1 with an Avitag. **C.** Metaplots and pause index of anti-Avitag eNETseq for WT and ΔCTD RNAPII. Note reduced RNAPII at the PPP relative to the gene body in ΔCTD (blue arrows). **D.** eNETseq for WT and ΔCTD RNAPII. Note reduced RNAPII at the PPP relative to the gene body (red arrows) and delayed termination (blue arrow) in ΔCTD. Anti-Avitag eNETseq in cells not expressing tagged RNAPII had a very low background (Fig. S3D). **E.** Metaplots of anti-Avitag eNETseq showing the 5’ shift in the PPP when the CTD is deleted (blue arrow).

## Discussion

We report a previously unrecognized dependency of RNAPII pausing on upstream G residues that was discovered by DEFT an AI that used an embedded LLM to discover DNA sequence features that distinguish between pause sites and non-pause sites. The decision tree approach employed by DEFT could in principle be used to find complex features that distinguish other functional classes of DNA sequences. We exploited the G-dependence of pausing to eliminate the PPP by insertion of a G-less ITS at two human genes. Ironically, G-less cassettes that prevent PPP formation in vivo were used in seminal studies of the PPP in vitro ^17,18^ where limiting nucleotide concentrations likely favored sequence-independent pausing. G-less insertion at the 5’ end is likely to be a general method for gene-specific PPP suppression. Inhibition of elongation by CDK9 inhibition with DRB could not restore the PPP after it had been suppressed by a G-less sequence (Fig. 3B, S1D). G-dependent pausing occurs throughout genes and can not explain why the PPP is restricted to 5’ ends, or why the ITS can no longer support an RNAPII peak if it is separated from the promoter by an insertion (Fig. 2C). We propose a multistage model of PPP establishment initiated by G-dependent arrest and subsequently stabilized by multiple factors including the +1 nucleosome^39^, NELF, DSIF ^15^ and promoter-tethering by the CTD (Fig. 7) ^44^. We found that the natural ITS of NDRG1 is not sufficient to elicit a PPP when separated from the promoter by a 380 base insertion (Fig. 2C, E). Therefore proximity to the start site appears necessary for PPP formation consistent with promoter-tethering that restricts RNAPII release within a limited distance from the TSS. We report that deletion of the CTD reduces PPP formation consistent with a previous report that CTD truncation reduces the duration of the PPP ^66^. CTD deletion also shifted the position of the PPP to a more upstream position (Fig. 6E, S3F) as predicted if this domain tethers RNAPII to the promoter by making long-range bridging contacts spanning ∼ 50-100 base pairs ^44^. Our work does not reveal how the G-richness of the nascent RNA or the non-template DNA strand promotes pausing, but interestingly R-loops and G-quadruplexes, enriched at the 5’ ends of human genes are favored by G’s and implicated in pausing ^67-69^. Our work does not reveal how the G-richness of the nascent RNA or the non-template DNA strand promotes pausing however a stable rG-dC base pair at the beginning of the transcription bubble is known to inhibit translocation by E coli RNA polymerase^70^. Furthermore R-loops and G-quadruplexes, enriched at the 5’ ends of human genes are favored by G’s and have been implicated in pausing ^67-69^.

**Figure 7.**
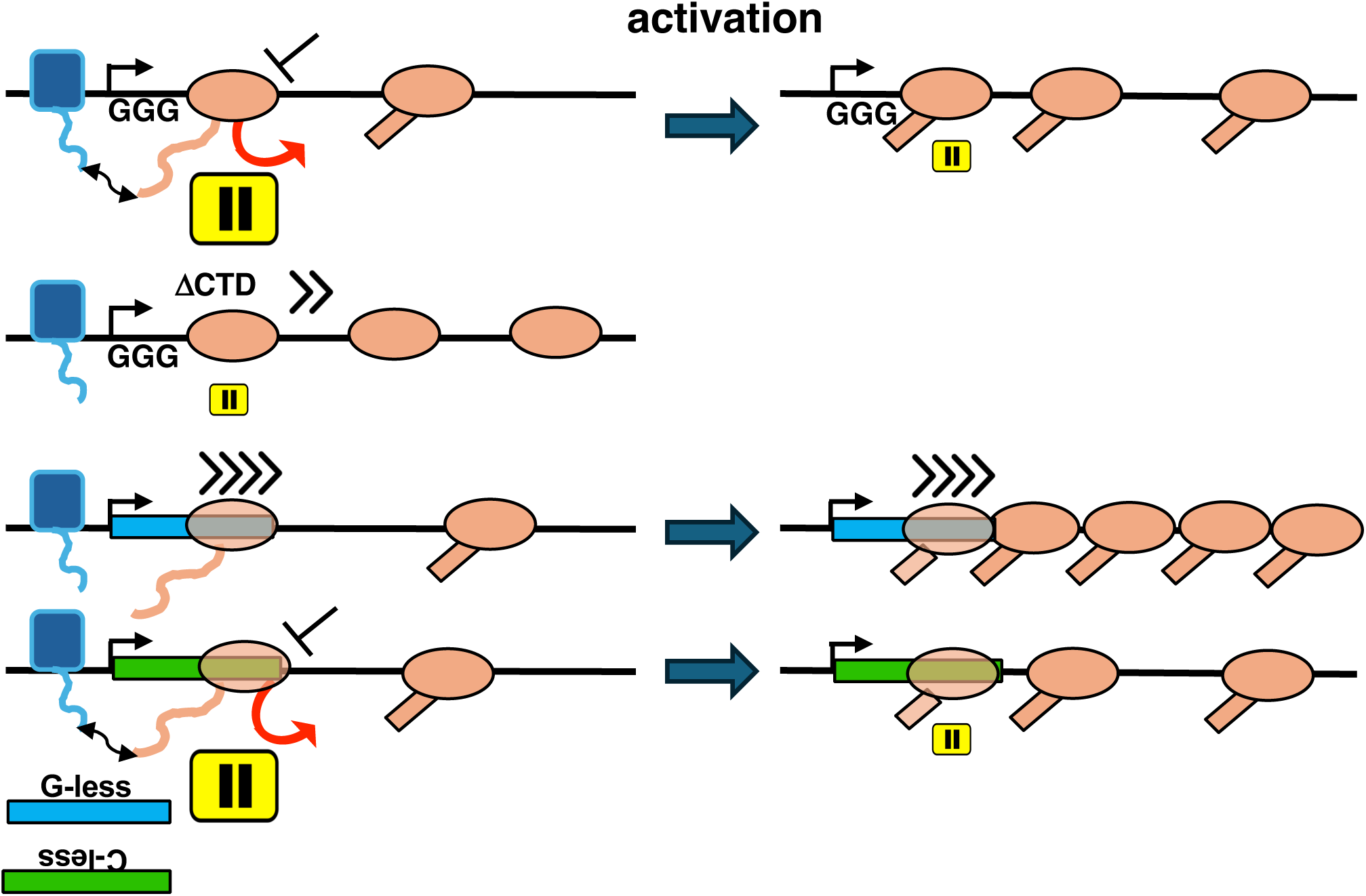
Model for G- and CTD-dependent PPP formation and gene activation independent of pause release. Red arrow indicates premature termination, double headed arrow indicates putative CTD-mediated tethering to promoter-bound transcription factors^43,44^.

Unexpectedly the abolition of the PPP by a G-less ITS, as detected by eNETseq, does not affect the ChIP-seq peaks of total RNAPII or Ser5-P RNAPII at the 5’ end of the gene (Fig. 3C, D, 5C). Thus, contrary to what is commonly assumed, the signature 5’ peak of RNAPII does not necessarily correspond to paused transcription elongation complexes. Instead we propose that the 5’ peak includes a large fraction of RNAPII at a pre-pausing stage of the transcription cycle where the nascent transcript is too short to be detected by eNETseq. We speculate that these early complexes correspond to polymerases that fail to clear the promoter and undergo abortive initiation which is very frequent in vitro ^34,35^ but currently impossible to assay in vivo. These results emphasize that the signature 5’ peak of RNAPII occupancy on metazoan genes should be interpreted cautiously. It is likely to include not only paused elongation complexes but also a large fraction of pre-pausing complexes that undergo rapid turnover.

Transcriptional activation through release of RNA polymerases from the PPP into productive elongation through gene to produce functional full length transcripts is an almost universally accepted model of eukaryotic gene regulation ^2,4-8,71^. However it has never been possible to test this model in a straightforward way by testing the effects of eliminating the PPP at a specific gene. We show here that suppression of the PPP at NDRG1 and HSP90AA1 by insertion of a G-less sequence at the 5’ end of the gene does not impair robust transcriptional activation in response to hypoxia or heat shock, contrary to the pause-release model ^2,4,8^. What then is the function of the PPP? One possibility is suggested by the high levels of RNAPII turnover by premature termination at the PPP ^27-29,72,73^. Stable pausing may be a pre-requisite for this turnover which likely functions in quality control and attenuation of RNAPII flux through genes ^3,6,31,74^. Reduced premature termination as a result of PPP suppression could explain both NDRG1 super-activation (Fig. 4A-C, E, S1E), and the reduced pausing index associated with CTD deletion (Fig. 6C, D, S3E). Finally we note a novel prospective role for AI in this study. Rather than just analysis of data, it discovered the feature, G-dependent pausing, that made a previously unconsidered causal experiment possible, PPP suppression by G-less insertion, and that experiment overturned the widely accepted relationship between promoter-proximal pausing and gene activation.

### Limitations of this study

The strategy of G-less cassette insertion to suppress the PPP was successful at the two highly regulated genes where we attempted it, however we do not know how general this method will be. It needs to be tested at other classes of promoter including CpG island promoters and those with diffuse start sites. The molecular basis for G-dependent pausing is unknown. Nor is it known whether G’s in the nascent transcript or on the non-template DNA strand are the signals for pausing. Further study will be required to identify the minimal sequence requirements for PPP formation and whether G-dependent R-loops and G-quadruplexes are involved. Although pause-release does not play a significant role in activation of a hypoxia-responsive gene and a heat-shock responsive gene, it is possible that there are other highly regulated genes for which it is required. We do not know the status of RNAPII complexes that are recruited to a promoter and give rise to large ChIP peaks, but are not paused elongation complexes that can be captured by NETseq. They may correspond to abortive initiation complexes seen frequently in vitro but it will be a challenge in future to establish whether this phenomenon is also common in vivo.

## Supporting information

DEFT metrics

Oligonucleotides

## Resource availability

Sequencing Datasets are deposited at GSE336158 and GSE336160. Other materials are available on request from.

## Acknowledgements

We thank U. Hyder, R. Treisman, D. Taatjes, and J. Svejstrup for helpful discussions, U. Hyder for technical help and the UC Denver sequencing facility. D.B. thanks the Francis Crick Institute. Supported by NIH grant R35GM118051 to D.B.

## Author contributions

D.B. and M.vdS. conceived the project. N.F. and D.B. designed experiments. N.F. performed all wet lab experiments and N.H. performed DEFT experiments. B.E. R.S. and N.F. performed bioinformatics. D.B. wrote the paper with input from all authors.

## Star Methods

### DEFT

DEFT model performance was assessed using a held-out set of 24,178 pause and 24,179 non-pause sites. Non-pause sites are positions drawn randomly from the same set of genes used to identify pause sites (longer than 2 kb and separated by at least 5 kb, the bottom 10% of genes with the lowest signal were excluded).

Definition of the metrics: Let TP, TN, FP and FN denote the numbers of true positives, true negatives, false positives and false negatives, respectively, with pauses defined as the positive class. We computed accuracy as (TP+TN)/(TP+TN+FP+FN). We additionally report the area under the receiver operating characteristic curve (AUROC), evaluated across classification thresholds (Table S1).

### Human cell lines

HCT116 cells were maintained in McCoy’s medium supplemented with 10% FBS and penicillin/streptomycin.

HAP1 cells ^75^ were grown in IMDM (GE Healthcare Bio-Sciences, Pittsburgh, PA) supplemented with 10% FBS, 4mM L-glutamine, and 1% pen/strep.

HEK293 Flp-in cells with integrated pcDNA5/FRT/TO-Rpb1 Am^r^ constructs were maintained in DMEM 10% FBS, 200 μg/mL hygromycin B, 6.5 μg/mL blasticidin, 1% pen/strep.

Cells were treated with DMOG (2mM, 24 hr) or CoCl_2_ (300 μM, 24 hr) to activate hypoxia responsive genes. Heat shock was at 42° for 30 min. immersed in a water bath. DRB treatment was 100μM for 2 hr.

### CRISPR-Cas9 cell line construction

G-less cassette insertions into HAP1 and HCT116 cells were made using the ALT-R CRISPR CAS9 system (IDT) for homology directed repair. crRNA (Table S2) + Alt-R^®^ CRISPR-Cas9 tracrRNA (IDT, catalog 1072532) were annealed and assembled into RNPs with S.p. Cas9 Nuclease V3 (IDT, catalog 1081058) according to the manufacturer’s instructions and electroporated with ALT-R HDR enhancer V2 into HAP1 or HCT116 cells using a Neon 100-μL Kit (Thermo Fisher, catalog N10025) on the Neon™ NxT Electroporation System (Thermo Fisher, catalog NEON1S). The G-less cassette from pML(C_2_AT) (377 bp) ^57^ was PCR amplified with primers (Table S2) that provided ∼100 bases of homology upstream and downstream of the insertion site and 1 μg was used for electroporation plus 1 μg plasmid encoding ATP1A1 gRNA to allow co-selection for ouabain resistance ^76^. Ouabain resistant colonies were screened by PCR and sequencing of PCR products to confirm accurate integration of the G-less and C-less cassettes. Two independent HAP1 and HCT116 cell lines were isolated with the G-less insertion (HAP1#10, #12, HCT116#10, #20) and two independent HAP1 lines (HAP1rev#14, rev#15) with the C-less insertion at NDRG1. One HAP1 line was isolated with the G-less insertion at HSP90AA1.

### HEK293 Flp-in cell lines

HEK293 Flp-in (Invitrogen) lines expressing Avitag Am^r^ Rpb1 WT and ΔCTD were constructed as described ^77^ by Flp recombinase mediated recombination. The N-terminal B10 epitope tag of pcDNA5/FRT/TO-B10-Rpb1 Am^r^ ^77^ was replaced with an Avitag. pcDNA5 Avi Rpb1Am^r^ ΔCTD deletes residues 1593-1961 including all 52 heptad repeats but retains the CTD linker region and the C-terminal 10 residues necessary for protein stability ^78^.

### Antibodies

Homemade rabbit anti-RNAPII pan CTD and anti-Avitag have been described ^21^.

### ChIP-seq

ChIP of human extracts and mapping to the hg38 UCSC human genome (Feb. 2009) with Bowtie2 version 2.3.2 has been described ^21^. 1Rabbit anti-pan RNAPII CTD, rabbit anti-NELFA and 3E8 rat anti Ser5-P monoclonal plus anti-rat secondary was used. PCR duplicates were removed using bbtools clumpify and adapters were trimmed using bbtools bbkuk version 39.01. Read coordinates were collapsed and centered using deeptools bamCoverage version 3.5.1. Pol II ChIP-seq, metaplots include genes longer than 1 kb, separated by >2 kb, containing at least one read in each region.

### eNET-seq

eNETseq, mapping and pause calling were performed as described ^21^ except that commercial (NEB M0608S) rather than homemade decapping enzyme was used. Datasets from G-less and C-less insertion lines were mapped to custom genomes based on hg38 containing the relevant insertion. HEK293Flp-in cells were treated with 2.0 μg/mL doxycycline for ∼24hr to induce expression of Avitag Rpb1WT or Rpb1ΔCTD before nuclei were harvested for eNETseq. Although these constructs have a resistance mutation, cells were not treated with α-amanitin. Micrococcal nuclease digested urea-washed chromatin from 4-6 15cm plate of cells was used for each IP with 40 μl of anti-CTD or anti-Avitag antiserum pre-bound to 50 μl of proteinA Dynabeads. Libraries were prepared with UMI’s using the QIAseq miRNA UDI kit (Qiagen 331505). Filtering was performed to remove reads that did not align within 5 kb of a protein coding gene, or that aligned to a snoRNA gene. eNET-seq datasets were down-sampled so that all libraries being compared had the same number of aligned and filtered reads. Pauses are defined as positions where eNETseq signal is 3 standard deviations above the mean for a window of 200 bases upstream and downstream with a minimum of 5 reads at the pause and 5 additional reads in the surrounding window ^21,37^. eNET-seq metaplots (Fig. 6C, S3E) include genes >2kb long, separated by >5kb in the top 90% for expression and read counts were equalized on a per gene basis by subsampling. To be included each gene had to contain at least one read in the TSS region -30:300 and body region +301:pA and be shared by all datasets. After clustering, a subset of 347 genes with strong 5’ peaks were filtered out. Nucleotide frequency was determined for the region +/-10 bp around pause sites separated by >30 bp for genes >2 kb long, separated by >5 kb are shown. and sequence logos (Figs. 5E, S3G) were generated using ggseqlogo ^79^.

**RT-PCR** of cDNA primed on total DNAse treated RNA with random hexamers using M-MLV RT has been described ^80^. Primers are provided in Table S2.

### Informatics

G-content in the first 100 bases of the transcript (Fig. 5D, E, S3A, B) was calculated using TSS coordinates derived from CAGE in HEK293 cells ^81^.

**Tables S1**

DEFT metrics

**Table S2**

Oligonucleotides used in this study

**Figures S1-S4**

