## Supplemental Figs S1-4 for "AI discovery of sequence rules for RNA polymerase II pausing revises the pause-release model of gene activation"

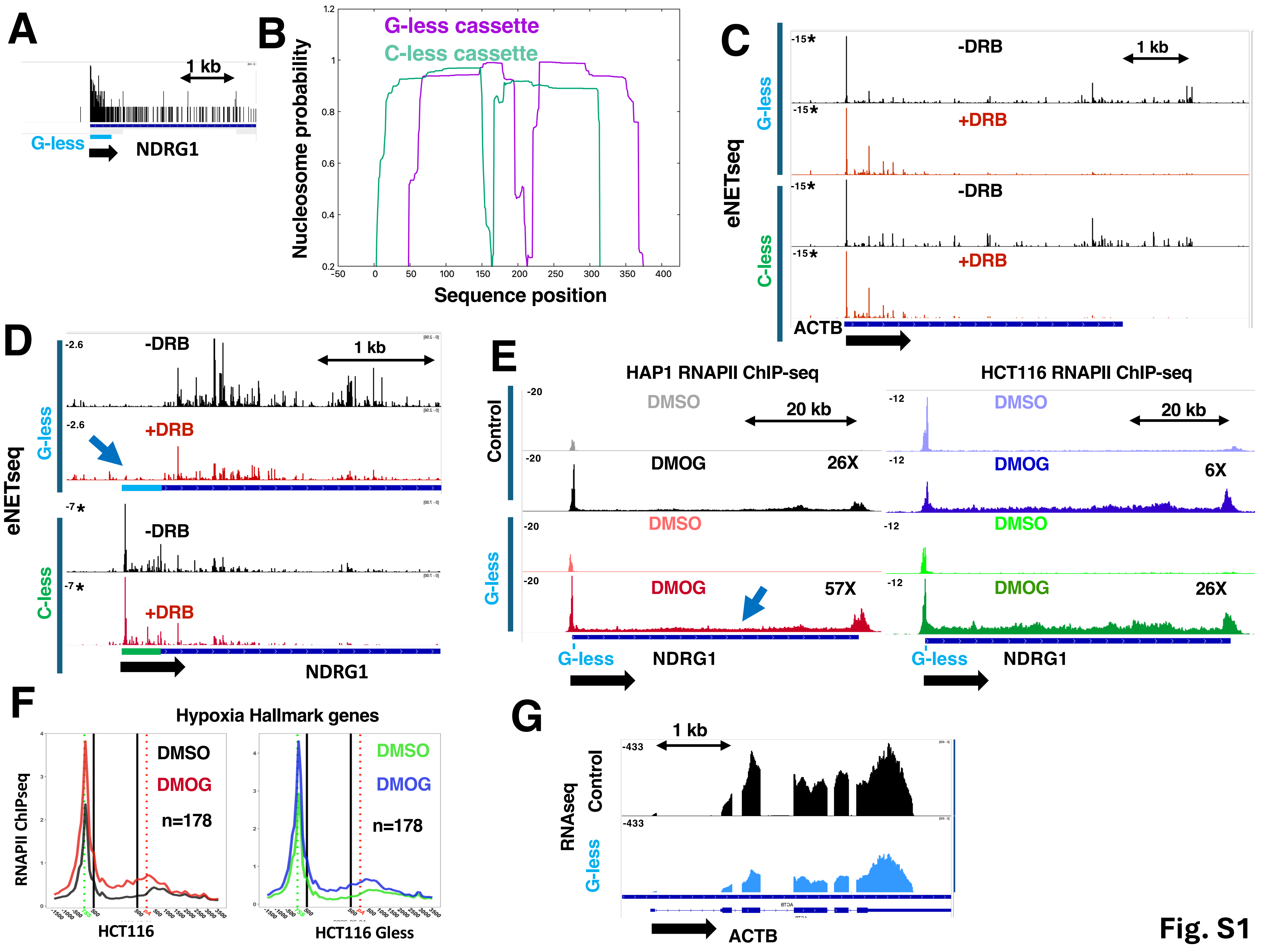
A
1 kb
G-less
NDRG1
B
G-less cassette
C-less cassette
Nucleosome probability
Sequence position
C
eNETseq
G-less
C-less
-15*
-DRB
+DRB
1 kb
ACTB
D
eNETseq
G-less
C-less
-2.6
-7*
-DRB
+DRB
1 kb
NDRG1
E
HAP1 RNAPII ChIP-seq
HCT116 RNAPII ChIP-seq
Control
G-less
-20
-12
DMSO
DMOG
20 kb
26X
6X
57X
26X
G-less
NDRG1
F
Hypoxia Hallmark genes
RNAPII ChIPseq
DMSO
DMOG
n=178
HCT116
HCT116 Gless
G
1 kb
RNAseq
Control
G-less
-433
ACTB

Fig. S1

**Figure S1**

**A.** Mapping of synthetic NETseq data to the NDRG1 gene with a G-less insertion. **B.** Predicted positions of two nucleosomes on the G-less and inverted C-less sequences using NuPop[61]. Note equivalent probabilities of nucleosome occupancy on the two sequences. **C.** eNETseq of ACTB in HAP1 with G-less and inverted C-less insertion at NDRG1 +/- DRB as in Fig. 3B. Note equivalent effects of DRB in both cell lines. * full range of the Y axis not shown. **D.** eNETseq of NRDRG1 with G-less and inverted C-less insertions +/- DRB (100 μM, 2 hr) in a replicate of Fig. 3B. * full range of the Y axis not shown. **E.** RNAPII ChIP-seq at NDRG1 1 in HAP1 and HCT116 cells -/+ G-less cassette insertion -/+ DMOG activation. Fold activation is based on gene body ChIP-signal. **F.** Metaplots of anti RNAPII ChIPseq for Hypoxia Hallmark genes (other than NDRG1) in control and NDRG1-G less insertion cells +/- DMOG. **G.** ACTB RNAseq in DMOG activated HAP1 cells (control for Fig. 4D).

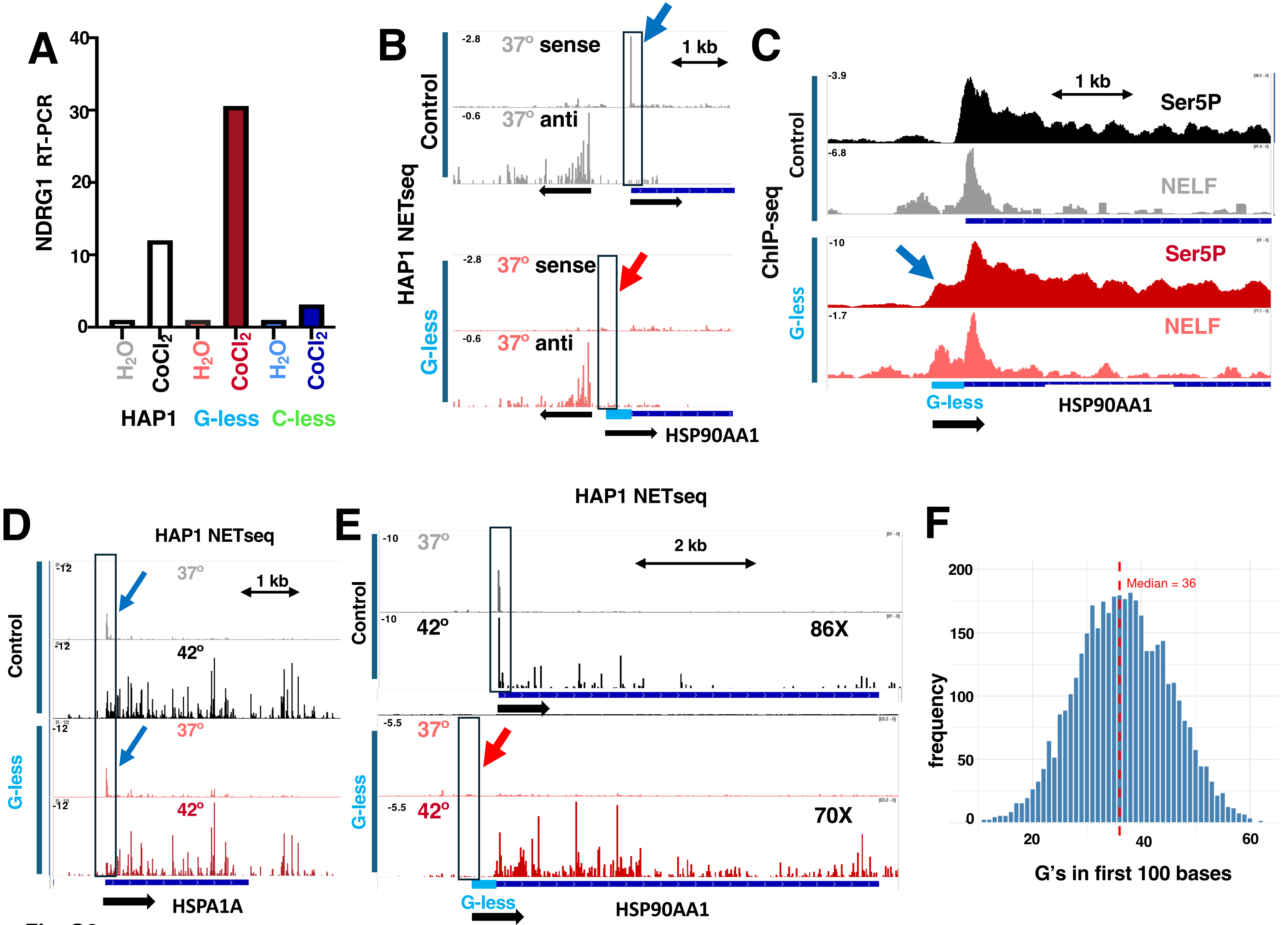
A
NDRG1 RT-PCR
40
30
20
10
0
H2O
CoCl2
H2O
CoCl2
H2O
CoCl2
HAP1
G-less
C-less
B
HAP1 NETseq
Control
-2.8
37° sense
1 kb
-0.6
37° anti
G-less
-2.8
37° sense
-0.6
37° anti
HSP90AA1
C
ChIP-seq
Control
-3.9
1 kb
Ser5P
-6.8
NELF
G-less
-10
Ser5P
-1.7
NELF
G-less
HSP90AA1
D
HAP1 NETseq
Control
-12
37°
1 kb
-12
42°
G-less
-12
37°
-12
42°
HSPA1A
E
HAP1 NETseq
Control
-10
37°
2 kb
-10
42°
86X
G-less
-5.5
37°
-5.5
42°
70X
G-less
HSP90AA1
F
200
150
100
50
0
frequency
Median = 36
20
40
60
G's in first 100 bases

Fig. S2

**Figure S2**

**A.** RT-PCR analysis of NDRG1 mRNA induction by $CoCl_2$ shows super-activation in HAP1 cells with the G-less cassette insertion but not with the inverted C-less insertion. **B. C.** Ser5-P and NELFA ChIP in HAP1 cells at 42º (30 min) -/+ Gless insertion at HSP90AA1. Note Ser5-P signal at 5' end of G-less cassette as expected for a TSS and the presence of NELFA in the absence of a PPP **D.** eNETseq at 37º and 42º (30 min.) at HSPA1 in HAP1 cells with and without a G-less cassette insertion at HSP90AA1. The PPP is marked (blue arrow). **E.** eNETseq at HSP90AA1 at 37º and 42º (30 min) in control HAP1 cells and HAP1 cells with G-less cassette insertion at HSP90AA1 as in Fig. 5B for a replicate experiment. Note the PPP present in control cells is abolished by the G-less insertion (red arrow). Fold activation of eNETseq reads in the gene body 42º/37º are marked. **F.** G content within the first 100 bases of the transcript for the top 50% of expressed genes based on RNAPII ChIP.

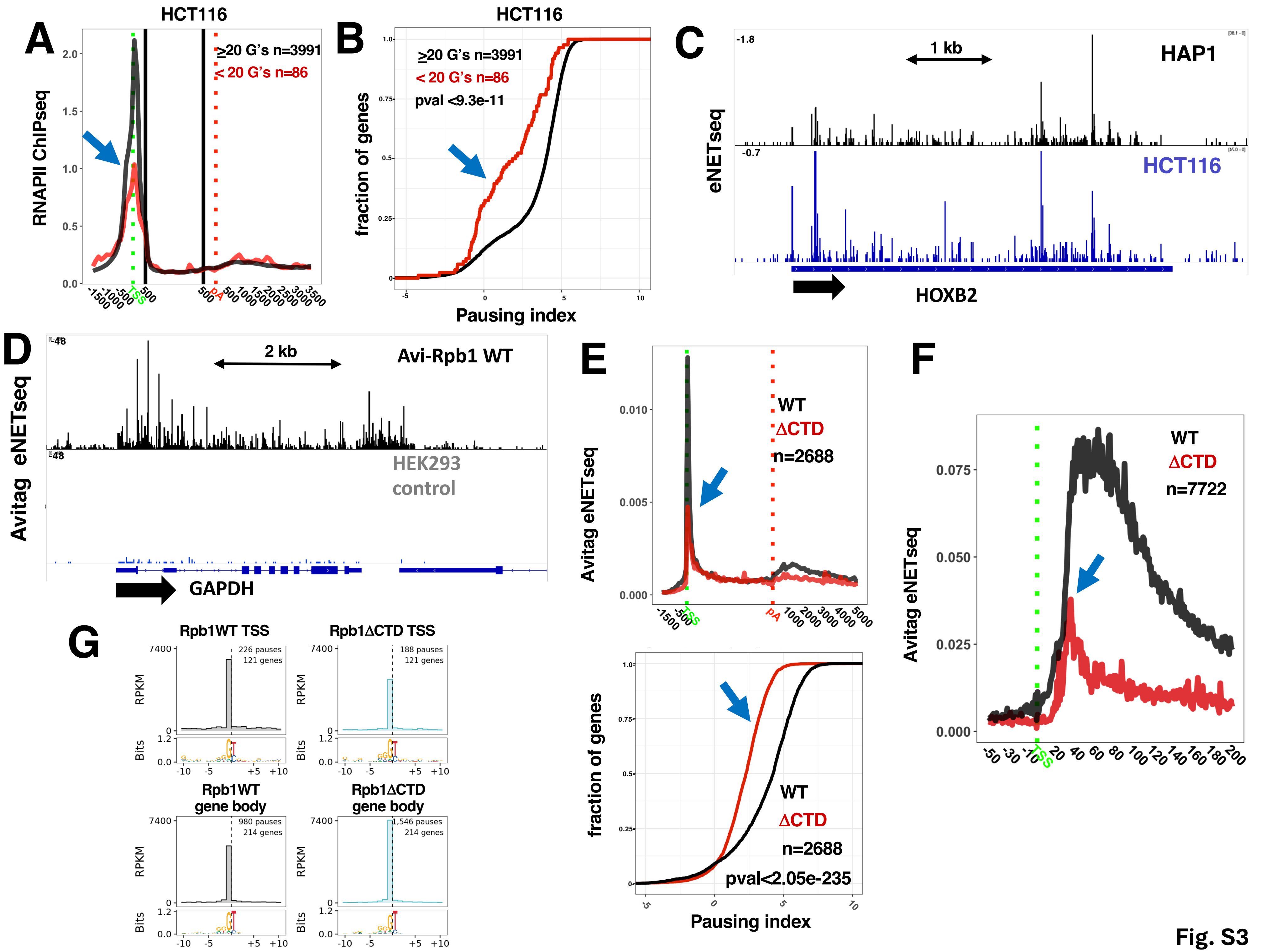
A
HCT116
RNAPII ChIPseq
≥20 G's n=3991
< 20 G's n=86
TSS
pA
B
HCT116
fraction of genes
≥20 G's n=3991
< 20 G's n=86
pval <9.3e-11
Pausing index
C
eNETseq
1 kb
HAP1
HCT116
HOXB2
D
Avitag eNETseq
2 kb
Avi-Rpb1 WT
HEK293 control
GAPDH
E
Avitag eNETseq
WT
ΔCTD
n=2688
fraction of genes
pval<2.05e-235
Pausing index
F
Avitag eNETseq
WT
ΔCTD
n=7722
G
Rpb1WT TSS
226 pauses
121 genes
Rpb1ΔCTD TSS
188 pauses
121 genes
Rpb1WT gene body
980 pauses
214 genes
Rpb1ΔCTD gene body
1,546 pauses
214 genes
RPKM
Bits

Fig. S3

**Figure S3**

**A.** RNAPII ChIPseq metaplots for genes in the top 50% for expression based with <20 or >20 G's in the first 100 bases of the transcript. **B.** Cumulative frequency plots of pausing index for genes with low and high G content as in A. Note that low G content is associated with significantly reduced promoter proximal pausing. **C.** eNETseq of HOXB2, a gene with low G-content in the ITS. Note the lack of a PPP. **D.** Anti Avitag eNETseq in control HEK293 Flp-in cells and cells expressing Avitag Rpb1 WT Am$^r$. Note low background signal in the control. **E.** Metaplot and pause index of anti-Avitag eNETseq results for WT and ΔCTD RNAPII as in Fig. 6B for a biological replicate Note reduced RNAPII at the PPP relative to the gene body as a result of CTD deletion. **F.** Metaplots of anti-Avitag eNETseq showing the 5' shift in the PPP when the CTD is deleted (blue arrow) as in Fig. 6D for a biological replicate. **G.** CTD deletion does not affect the sequence preference at RNAPII pause sites. Consensus sequences around the pause site (dotted line) for WT and ΔCTD RNAPII as in Figure 5E for replicate eNETseq experiment.

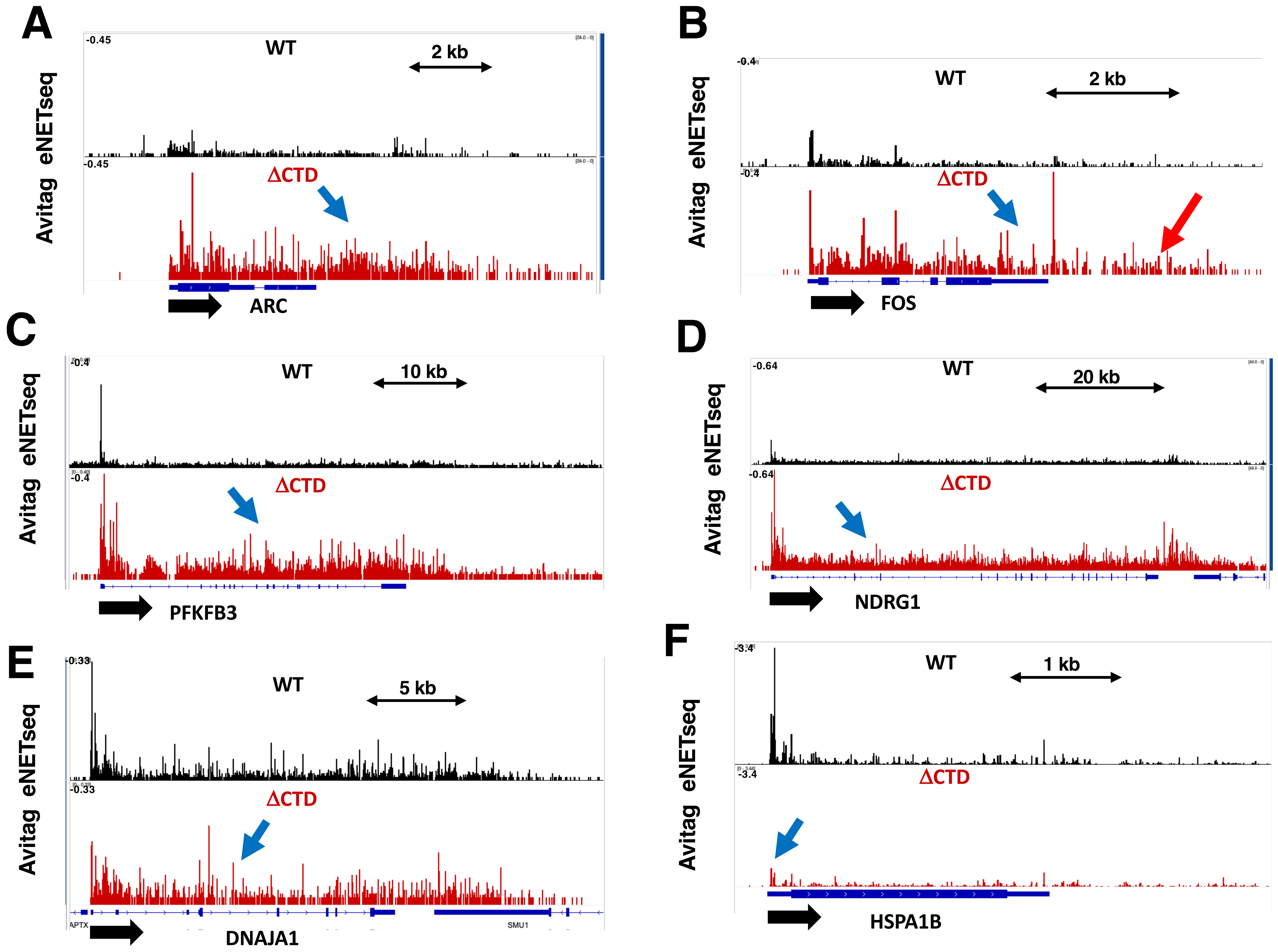
A
Avitag eNETseq
-0.45
WT
2 kb
-0.45
ΔCTD
ARC
B
Avitag eNETseq
-0.4
WT
2 kb
-0.4
ΔCTD
FOS
C
Avitag eNETseq
-0.4
WT
10 kb
-0.4
ΔCTD
PFKFB3
D
Avitag eNETseq
-0.64
WT
20 kb
-0.64
ΔCTD
NDRG1
E
Avitag eNETseq
-0.33
WT
5 kb
-0.33
ΔCTD
APTX
SMU1
DNAJA1
F
Avitag eNETseq
-3.4
WT
1 kb
-3.4
ΔCTD
HSPA1B

Fig. S4

**Figure S4**

**A-F.** Screenshots of inducible genes anti Avitag eNETseq in HEK293 Flp-in cells expressing Avitag-Rpb1 WT or ΔCTD. Note suppression of the PPP relative to RNAPII occupancy in the gene body.
